# Neurons as biosensors for discriminating neurological disorders in a brain-on-chip platform: Application to Alzheimer’s Disease using patient CSF

**DOI:** 10.1101/2024.08.23.609425

**Authors:** Louise Miny, Jessica Rontard, Ahmad Allouche, Nicolas Violle, Louise Dubuisson, Aurélie Batut, Alexandre Ponomarenko, Rania Talbi, Hélène Gautier, Benoît Maisonneuve, Serge Roux, Florian Larramendy, Thibault Honegger, Isabelle Quadrio

## Abstract

Alzheimer’s disease (AD) is characterized by the accumulation of aggregated amyloid beta peptide (Aβ) leading to progressive neuronal loss and dysfunction. Current AD’s diagnosis involves biomarkers assays in cerebrospinal fluid (CSF) as Aβ to validate the diagnosis. However, these methods are time-consuming, expensive, and can result in inaccurate diagnoses by not accounting for differential diagnose. To overcome these challenges, researchers are exploring new technologies for detecting AD biomarkers in biological fluids, though progress is hindered by an incomplete understanding of AD mechanisms and CSF composition. In this study, we used a standardized microfluidic platform to investigate the effects of synthetic Aβ peptides and cerebrospinal fluid (CSF) from AD and healthy patients on neuronal functional activity. First, human neurons derived from induced pluripotent stem cells (iPSCs) were characterized. Then, to modulate the functional activity of neurons, tetrodotoxin (TTX), a specific sodium channel blocker, was used as a control for inhibiting neuronal activity. Subsequently, glutamatergic neurons were chronically exposed to AβO and patients’ CSF. MEA recordings were performed before and after the treatments to assess changes in network activity. Our results demonstrated that extracting key electrophysiological metrics allows for discrimination between healthy and AD CSF samples. This system could offer the potential for differential diagnosis and development of personalized therapeutic strategies.

## Introduction

Alzheimer’s disease (AD) is the first cause of dementia in the world. One of its major hallmarks is the accumulation of aggreged amyloid-beta peptide (Aβ) in neurons leading to synaptic dysfunction and neuronal death and has been commonly studied for decades ^1–4^. Currently, the diagnosis is based on a combination of clinical examination on the patient’s symptoms, neuropsychological testing, and brain imaging. In atypical presentation, core biomarkers of AD neuropathological changes can be used to establish the diagnosis and precise the etiology. Indeed, among fluid biomarkers, cerebrospinal fluid (CSF) Aβ measurement, reflects the amyloid-beta pathway and helps to valid or invalid AD diagnosis ^5–7^. Unfortunately, these processes are long, expensive, and could lead to inaccurate diagnoses because differential diagnoses are not considered. This is a real challenge for public health and for the quality of life of patients and their families ^8^.

Since the early 2000s, researchers have attempted to develop innovative and reliable technologies to improve the detection of AD biomarkers in biological fluids ^9–12^. However, the incomplete understanding of AD mechanisms and the precise composition of CSF in AD patients makes it challenging to develop accurate diagnostic tools and effective treatments. These factors are essential for ongoing research focused on elucidating the pathophysiology of AD and thoroughly characterizing CSF biomarkers.

Microfluidic technology has been used for decades to create devices for the study and detection of several neurological diseases using biological fluid ^13–15^. For example, Koch et al, in 2019, used a microfluidic device coupled with microelectrode array (MEA) where neuronal culture could be recorded, allowing to discriminate between CSF from healthy and autoimmune encephalitis (AE) patients.

Electrophysiological recordings using MEA technology, are current readouts in neurological in vitro studies to evaluate the maturity, functionality and robustness of neuronal networks in microfluidic devices ^16–19^. However, most of published research so far has been using microfluidic devices coupled with MEA technology in single-device recording ^20–25^.

In microfluidic devices with compartmentalized neuronal cultures, neurons could act as sensors by responding to stimuli or drugs added to a specific channel. This setup allows precise control the microenvironment of linked neuronal populations, enabling researchers to observe how a drug applied in one compartment affects the entire interconnected network. By monitoring changes in neuronal activity across compartments, this approach could provide insights into the drug’s impact on neuronal activity and network dynamics.

In this study, we aimed to prove that neurons can be used as a biosensor and the neural network activity can discriminate pathologies in a high throughput (HTS) organ-on-chip platform. First, glutamatergic and GABAergic neurons from human induced pluripotent stem cells (hiPSCs) were seeded in compartmentalized high throughput devices. Second, we measured the reliability and reproducibility of the in vitro compartmentalized cell culture using electrophysiological recordings of neurons functionality, with TTX, an ionic channel inhibitor, and synthetic Aβ oligomers. Finally, CSF from healthy and AD patients was added to human glutamatergic neurons and showed that the extraction of key neuronal network activity metrics can be used to discriminate healthy or AD patients CSF.

## Results

### An Organ on chip platform for high-throughput electrophysiological recording of compartmentalized neural networks

To simultaneously record neurons co-cultures, a microfluidic platform composed of 16 microfluidic devices coupled with MEA in a standard high throughput design, named DuaLink MEA, was developed and manufactured (Fig. 1a). Microfluidic devices are made of Polydimethylsiloxane (PDMS) (Fig. 1b). Devices are composed of inlets and outlets, to seed cells and adding compounds, and three compartments linked with microchannels allowing fluidic isolation. The dimensions of both cell culture channels are 18800 (Length) × 1000 (Weight) x 200 (Height) µm, and microchannels are 125 (Length) x 6 (Weight) x 3 (Height) µm (Figure S1)). This architecture allows neurons to be seeded in the cell culture compartment (channel 1) while only neurites can grow throughout microchannels to reach the opposite cell culture (in channel 3) (Fig. 1c). Moreover, microfluidic devices were directly bonded on a MEA sheet developed by Axion Biosystems allowing neurons to be in direct contact with electrodes. (Fig. 1c). All regions of the neuronal network were recorded: 15 electrodes were placed in both channels 1 and 3 (Ch1 and Ch3), 5 electrodes in the inter-microchannels (1μ2 and 2μ3), and 2 electrodes in channel 2 (Ch2) (Fig. 1c). For easier interpretation of the results presented in this paper, the electrodes were color-coded according to their position under the device (each compartment/set of microchannels having its specific color, as depicted in Fig1d. Compartmentalization allowed us to investigate functional activity across neurons and neuronal network (Fig 1d). This specific positioning of electrodes allowed us to record various neuronal components, including neuronal culture in channels as well as their axons and dendrites in microchannels. Microchannels specifically enabled the isolation of axons and neurites and the precise recording of action potential between channels. Thanks to the transparency of the PDMS and the MEA sheet, the neuronal network organization and composition were investigated using immunofluorescence assay as exemplified in supplementary Figure S2.

**Figure 1:**
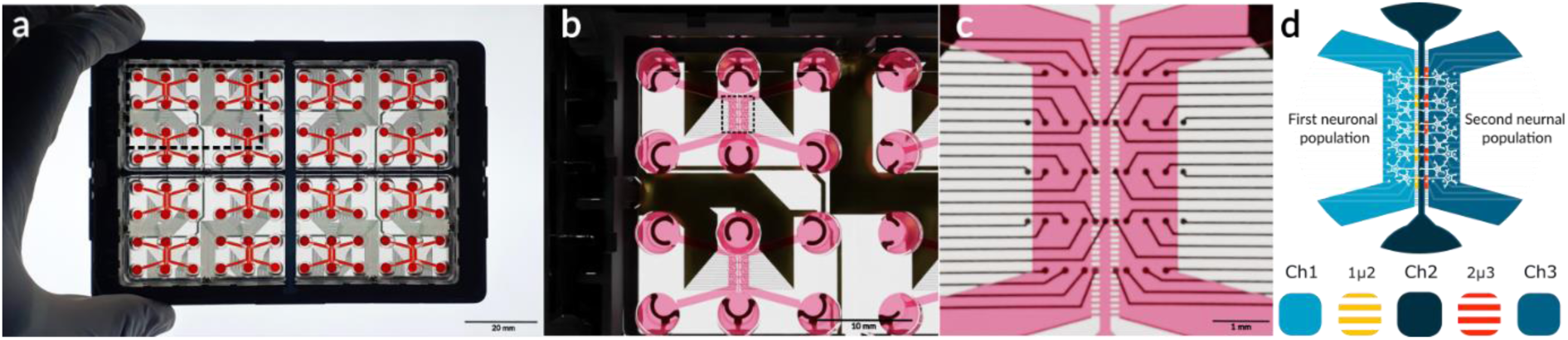
Microfluidic platform by NETRI with Axion Biosystem’s technology. (a) Pictures of the NeoBento Pharma composed of four QuarterBento and sixteen microfluidic chips. (b) Rendering of zoomed QuarterBento of four DuaLink chip MEA. (c) Realistic rendering of a detailed DuaLink chip MEA showing three culture compartments. Black dots represent microelectrodes placed below the chip. (d) Schematic representation of DuaLink MEA with compartmentalized neuronal culture with one cell type seeded in Ch1 and the other cell type seeded in Ch3, axonal propagations are respectively in microchannels 1µ2 and 2µ3.

### Neurons hiPSC derived in chip are mature and present spontaneous activity

Human glutamatergic and GABAergic neurons derived from iPSCs were seeded in DuaLink MEA’s Ch1 and Ch3, respectively. Using adapted provider protocols to ensure full differentiation and maturation prior to performing subsequent experiments, they were maintained for 21 days and supplemented with fresh media to achieve fully maturation between day 19 and day 21. Brightfield pictures were taken to monitor the evolution of neurons and their connection over time (Fig. 2b). Neuronal structure and functional activity were confirmed with immunofluorescence assay at day 21 and electrophysiological recordings of spontaneous activity at days 7, 14, and 21. Moreover, neuronal viability in microfluidic device were evaluated at the end of the culture with Live/dead assays with quantification (Fig. 2a). As expected from the cells’ providers’ protocols, the viability of the glutamatergic and GABAergic neurons at day 21 was 46.7 (±8.4) % and 49.5 (±10.2) %, respectively (n=20) (Fig. 2c). There was no significative difference of the live cells’ percentage between glutamatergic and GABAergic neurons (Fig 2f). To ensure the full differentiation of iPSC in neurons, pluripotency and specific markers were quantified by immunofluorescence at day 21 (n=10 per markers) (Fig. 2d and e). Pluripotency markers (Sox2 and Nestin) stained respectively 4.7 (±4.9) % and 1.5 (±2.4) % of the glutamatergic neurons, and 9.7 (±12.6) % and 4.2 (±6.5) % of the GABAergic neurons. They were both inferior to 10%, indicating that cells were differentiated (Fig. 2g). β-III tubulin, a specific neuronal marker, stained 85.2 (±6.7) % of the glutamatergic population and 78.1 (±19.3) % of the GABAergic population. For both neuronal types, vGlut1 (glutamatergic specific marker) and GABA (GABAergic specific marker) stained respectively 84.3 (±18.1) % and 3.8 (±4.1) % of glutamatergic neurons, and 2.1 (±2.9) % and 69.4 (±3.8) % of GABAergic neurons (Fig. 2h). We confirmed that most of the neurons presumed to be glutamatergic are indeed glutamatergic, and those presumed to be GABAergic are indeed GABAergic. Similarly, neurons presumed to be GABAergic were verified by the presence of the GABA marker, confirming their identity as GABAergic neurons. Finally, the neuronal co-culture recorded at day 7, 14 and 21 allows to plot raster graphs showing a global observation of the total recording over time on all electrodes. They showed an increase of detected action potential, represented by compartments color-coded points, over time (Fig. 2i). The recorded activity in the channels over time, represented by weighted mean firing rate (WMFR, i.e. number of spikes per seconds on active electrodes), showed a tendency to increase between day 7 and day 14. WMFR becomes statistically significant in microchannels between day 14 and day 21 (p<0.0001). At day 21, WMFR was increased in microchannels compared to channels (p<0.0001) (Fig. 2j). Synchrony of both glutamatergic and GABAergic neurons was also measured. A significant rise of synchrony index, generally comprised between 0 and 1, was observed between day 14 and day 21 in microchannels and in the channel 3 (Fig. 2k).

**Figure 2:**
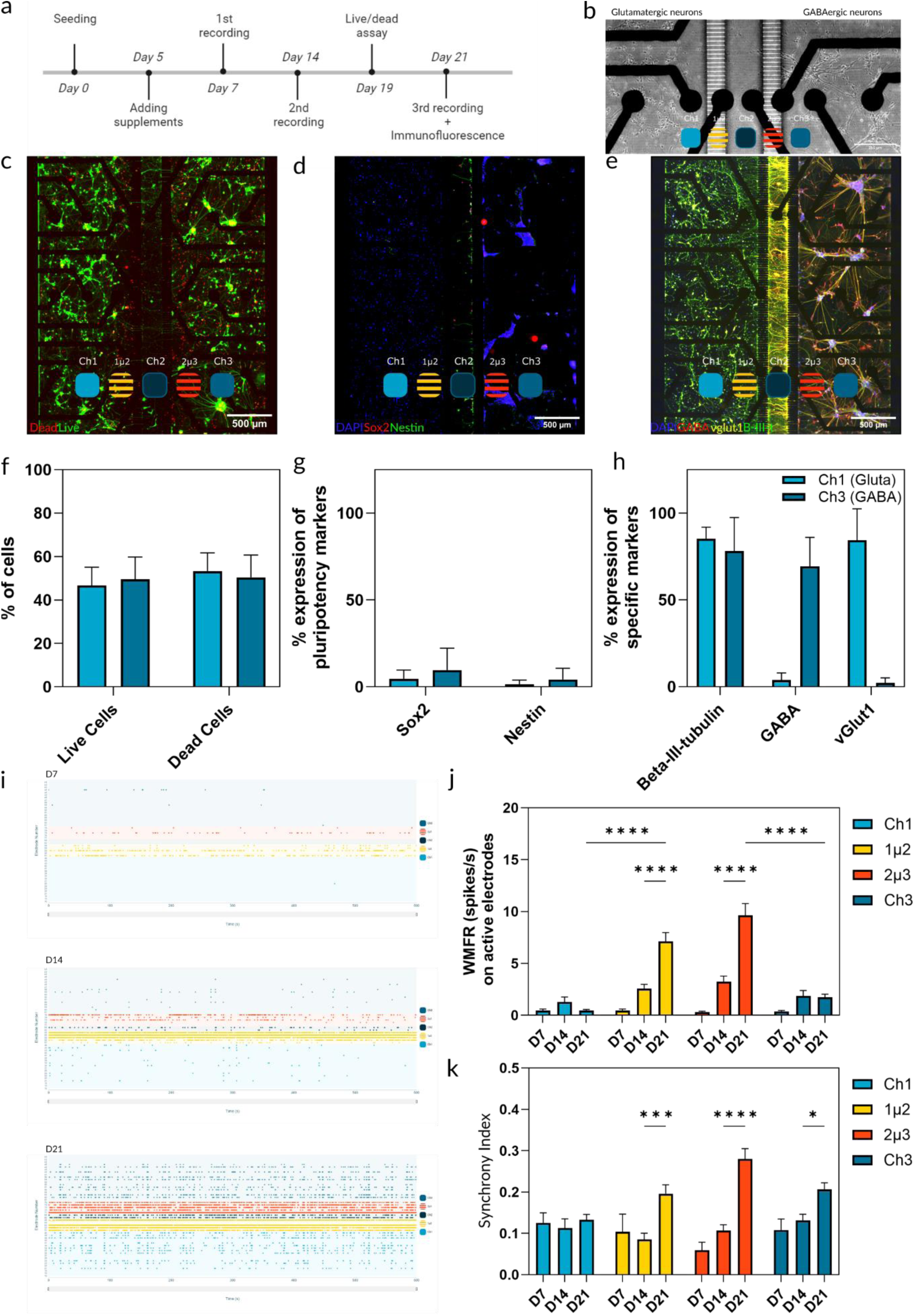
Characterization of co-culture of human Glutamatergic and GABAergic neurons in microfluidic device. (a) Timeline of cell culture maintenance with milestones during culture. (b) Brightfield contrast pictures of human glutamatergic and GABAergic neurons seeded in Dualink MEA at day 16. (c) Fluorescent picture of Live/Dead assay in the co-culture. Live cells were stained in green, and dead cells in red. (d, e) Immunofluorescent pictures of respectively, pluripotency markers (d), and specific neuronal markers (e) for each cell type. Sox2 was marked in red and Nestin in green. Β-III-tubulin was marked in green, vGlut1 in yellow, and GABA in red. In each immunofluorescent picture, DAPI was in blue. (f, g, h) Graph bars showing the quantification in channel 1 and channel 3 of the Live/dead assay, pluripotency markers and neuronal specific markers, respectively. (i) Raster plots from electrophysiological recordings at day 7, day 14 and day 21, showing spikes in time per electrode, in each DuaLink compartment. (j, k) Bar graphs showing, respectively weighted mean firing rate (WFR) and Synchrony index at day 7, day 14, and day 21 measured in channels and microchannels. * p-value< 0.05, *** p-value < 0.001 and **** p-value < 0.001. Error bar = SD.

### Chemical inhibitor placed in one compartment only have an impact over the electrical activity of the entire network

For all further experiments, we have used the DuaLink MEA in which glutamatergic neurons were seeded in both Ch1 and Ch3, while neurites could go throughout 1µ2 and 2µ3 respectively (Fig. 2b). By using a the same neuronal culture in both channels, we could assess with more accuracy the drug’s impact on synaptic connectivity and network activity, isolating the effects on excitatory transmission without the influence of inhibitory signalling from GABAergic neurons. This approach allowed us to gain clearer insights into the drug’s mechanism of action within a controlled excitatory neural network.

To ensure that neuronal co-culture could respond to chemical or biological perturbation, tetrodotoxin (TTX), a sodium receptor blocker well known to block synaptic transmission, was applied on neurons at a concentration of 10nM. Prior to TTX exposure, basal recordings were performed and showed spontaneous activity in both cultures. We added TTX at different time points to test the fluidic isolation and modulation of neuronal networks in separate compartments of our microfluidic devices. Specifically, we introduced TTX into one compartment (Ch1) on D18 before the experiments. Subsequently, TTX was administered to the second compartment (Ch3) at D19. It allowed sufficient time for the neurons in Ch1 to return to baseline functional activity, ensuring that any observed effects were localized and compartment specific. TTX effects on both glutamatergic neurons culture was recorded few minutes after the addition (Fig. 3a,b). At D18 and D19, raster plots showed a distinct diminution of detected spikes in the TTX-treated channel, and no change in functional activity in the opposite channel (Fig. 3c). The effect of TTX was compared to vehicle and showed a significative difference in percentage of change of WMFR and synchrony index to baseline. At D18, in Ch1 we observed a decrease of 97.7 (±4.5) % of change to the baseline of WMFR versus 11.5 (±7.5) % with vehicle. When TTX was added in Ch3, we observed a decrease of 76.3 (±53.0) % compared to 19.6 (±14.3) % with vehicle (Supplementary Fig. S3). In Ch1, the WMFR showed a tendency to decline within few minutes of TTX application between D18 and D18 (TTX Ch1) (ns) and no significative difference between D19 and D19 (TTX Ch3). However, we showed a significative decrease of WMFR in TTX-treated Ch3 between D19 and D19 (TTX Ch3) (p<0.05). Moreover, we observed a significative increase of WMFR between D19 (TTX Ch3) and D20 (p<0.05), showing a return to basal activity at D20 (Fig. 3 d). TTX was briefly applied in Ch1 on D18, resulting in a significant decrease in the weighted mean firing rate (WMFR) in 1µ2 (p<0.01) and 2µ3 (p<0.05). Similarly, applying TTX to Ch3 on D19 led to a significant reduction in WMFR in 2µ3 (p<0.001) and 1µ2 (p<0.05). We hypothesize that neurites from glutamatergic neurons in Ch1 extended into Ch3 through the 1µ2 and 2µ3 compartments and neurites from glutamatergic neurons in Ch3 extended into Ch1. This connectivity could explain the observed decreases in activity in the 2µ3 region when TTX was applied in Ch1, and in the 1µ2 region when Ch3 was treated with TTX (Fig. 3e). No significant difference in WMFR between D18 and D20 in channels nor microchannels was observed, showing the reversible effect of TTX on glutamatergic neurons (Fig. 3d,e). We have also observed that electrophysiological detection is improved in microchannels compared to channels. Indeed, basal activity in microchannels was about 46 spikes/s (Fig. 3d) on active electrodes while it was about 0.6 spikes/s in channels (Fig. 3c). Considering these results, we choose to only present microchannels (1µ2 and 2µ3) data for the following experiments. Furthermore, only the microfluidic devices that showed a successful response to TTX treatment, with evidenced by a significant decrease in neuronal activity, were selected for subsequent exposure to Aβ oligomers and CSF. This selection criterion ensured that the devices used in the Aβ and CSF experiments had functional networks capable of demonstrating the expected pharmacological effects.

**Figure 3:**
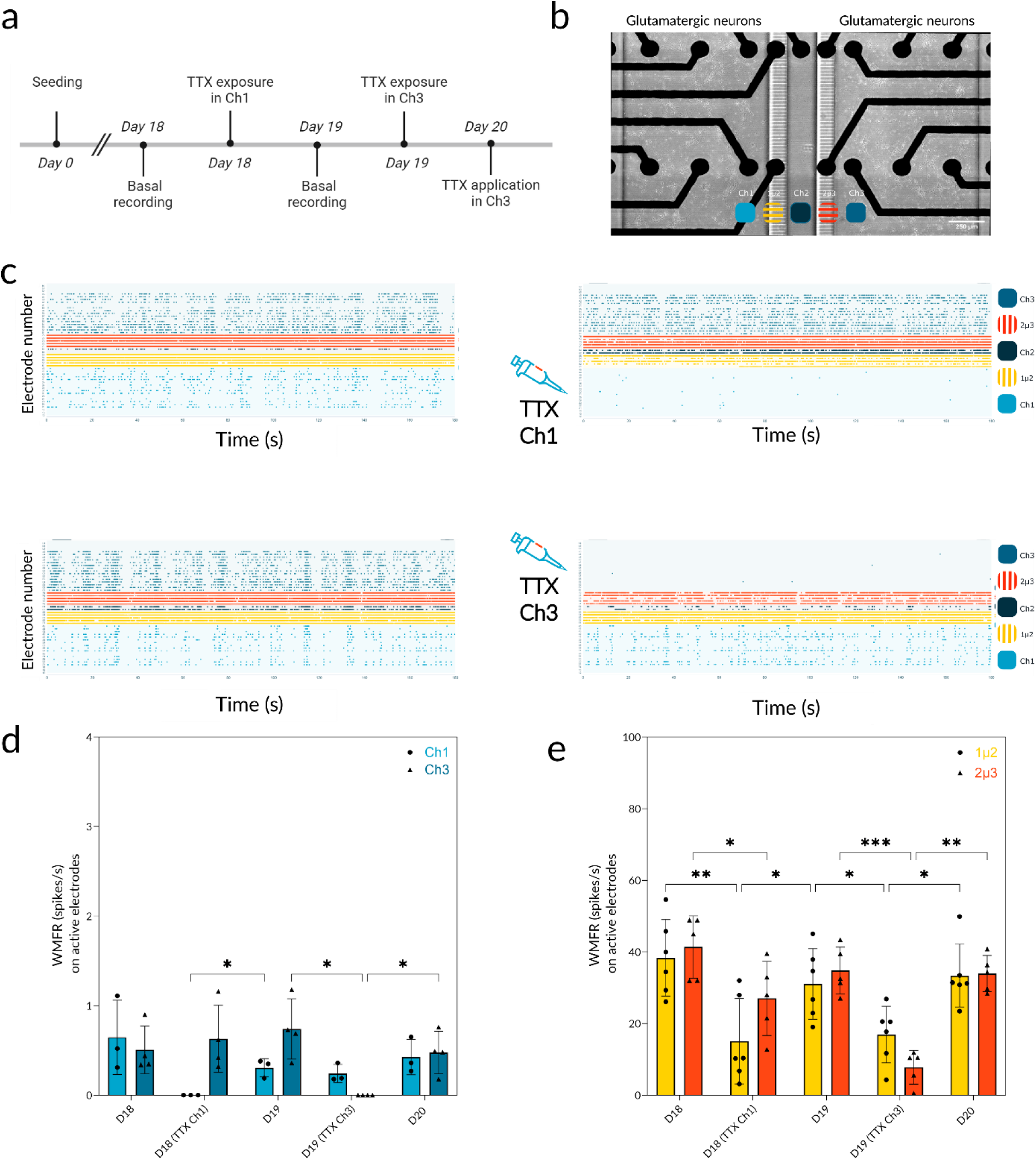
Electrophysiological modulation of co-culture glutamatergic neurons with TTX. (a) Timeline of TTX exposure experiment steps. (b) Brightfield contrast pictures of compartmentalized glutamatergic neurons culture in Ch1 and Ch3, with the color legend of each compartment. (c) Raster plots before/after neuronal exposure of TTX in channel 1 or in channel 3. (d, e) Bar graphs showing WMFR in each step of TTX exposure in channels and in microchannels respectively. * p-value < 0.05, ** p-value <0.01, *** p-value < 0.001 and **** p-value < 0.001. Error bar = SD. TTX: tetrodotoxin. WMFR: weighted mean firing rate.

### Aβ oligomers peptides placed in one compartment only have an acute functional effect on the entire network

Aβ oligomers (AβO) were used at a concentration of 10 µM and was added in Ch1. To determine the composition of Aβ oligomers, we analysed them using a Coomassie Blue staining method in conjunction with gel electrophoresis. This approach allowed us to visualize the protein bands and assess the size distribution and aggregation state of the oligomers. By examining the gel profile, we could identify the different oligomeric forms present, providing detailed insight into the molecular composition of the Aβ samples used in our experiments (Supplementary Fig. S4). Basal recording was performed at D19 followed by chronic applications of AβO every 24h (Fig. 4a). The AβO vehicle was used as a control, following the same chronic application protocol. AβO exposure on glutamatergic neurons induced a decrease in recorded functional activity, that becomes significant after 72h of exposure in 1µ2 and 2µ3 (n=6, p<0.05) (Fig. 4b). In opposite, adding AβO vehicle on glutamatergic neurons didn’t affect the functional activity (n= 2, ns) (Supplementary Fig. S4).

**Figure 4:**
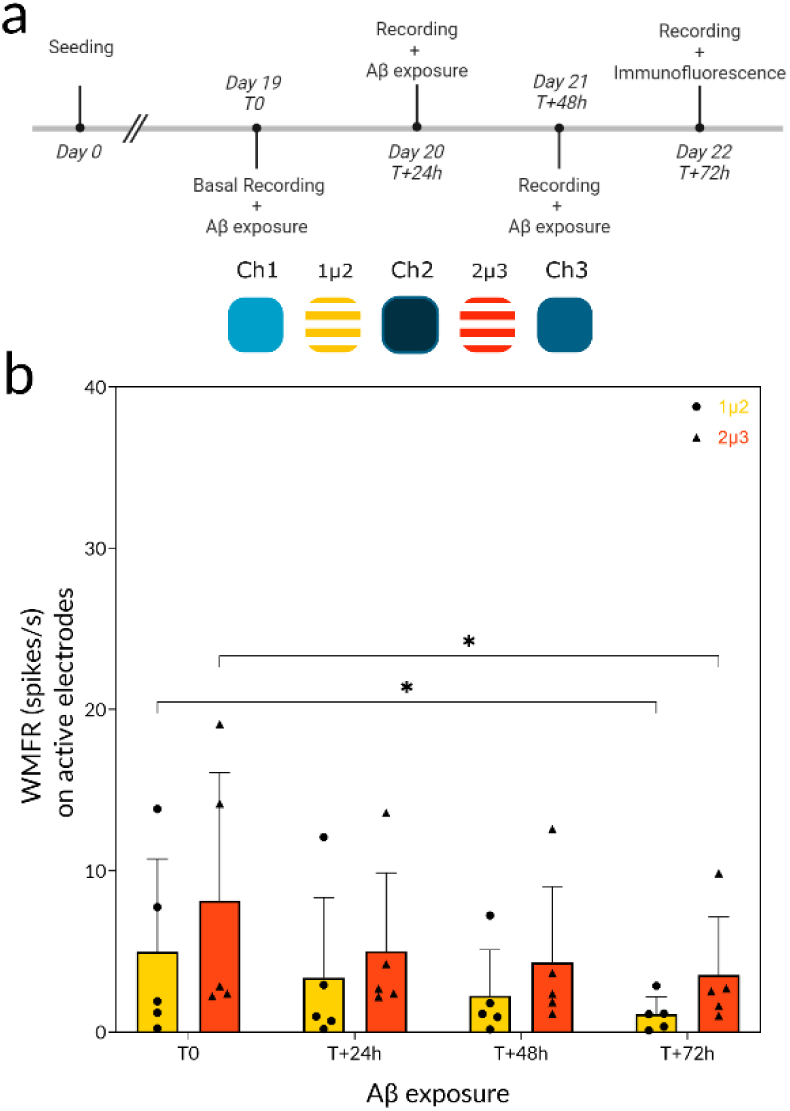
Electrophysiological effect of Aβ exposure on co-culture glutamatergic neurons. (a) Timeline of Aβ exposure experiment steps. (b) Bar graph of WMFR in microchannels of chronic Aβ exposure in time respectively. * p-value < 0.05, Error bar = SD. WMFR: weighted mean firing rate.

#### A. Effect of healthy and AD patients’ CSF exposure in neuronal network

Prior to experiments, CSF of healthy and Alzheimer’s patients were analysed, and biomarkers such as Aβ1-42, Tau total and p-281-tau were measured. Levels of Aβ1-42 tended to be lower in healthy patients compared to AD’s patients. There was no correlation between the tau level in CSF’s patients and the presence of the disease (Supplementary Table S1). Following the protocol described previously for the Aβ experiments, CSF from healthy (n=2) and Alzheimer’s patients (n=3) were chronically added on glutamatergic neurons in Ch1, at D19 every 24h during 72h (Fig. 5a). Initial exposure of AD CSF tended to increase neuronal activity at 24h in 1µ2 (p<0.01 for A1 and A3) and in 2µ3 (p<0.05 for A1 and p<0.01 for A2) (Fig. 5b, c, d). From 24h to 72h, no significant change of activity was observed. Healthy CSF from H1 patient didn’t change activity overtime (Fig. 5e). Surprisingly, the CSF of H2 patient tend to improve neuronal activity in 2µ3 compartment after 24h and 72h of exposure (Fig. 5f). The same experiment was performed with fresh media as a control. As expected because of the media composition, WMFR improved overtime in 1µ2 at 24h and 48h (p<0.05 and p<0.01) and in 2µ3 at 48h and 72h (p<0.05 and p<0.01) (Fig. 5g). To extend our analysis, we have explored the correlation between two electrophysiological metrics, with correlation matrix. All electrophysiological metrics has been used. Correlation matric shown that there was no strong correlation between WMFR and synchrony index highlighted for AD’s patients (Fig. 6a). We set a threshold of 0.7 to identify and exclude metrics with a strong correlation. Metrics that exceed this threshold were removed from the analysis to avoid overemphasizing any single metric, especially before performing a Principal Component Analysis (PCA). This step is crucial to prevent the inclusion of redundant information, which could otherwise lead to an imbalance in the analysis. We could choose the appropriate metrics to further analysis (in blue) (Fig. 6b). Scatter plots, showing the % of change to the baseline after 24h, 48h and 72h of exposure in 1µ2 microfluidic compartments were created. At the 24h the scatter plot shows a relatively mixed distribution of patient groups. Data points may be closely clustered, indicating that at this early stage, the differences between the patient groups are not yet pronounced. At 48h, the scatter plot begins to show a tendency for patient groups to differentiate. Data points may start to spread out, with some groups forming distinct clusters. Then, at 72h, the scatter plot likely reveals a more pronounced isolation and expansion of patient groups. The clusters representing different groups may be distinctly separated, with minimal overlap, indicating a significant differentiation based on the measured variables. This isolation suggests that each group is characterized by unique profiles or responses (Fig. 6c).

**Figure 5.**
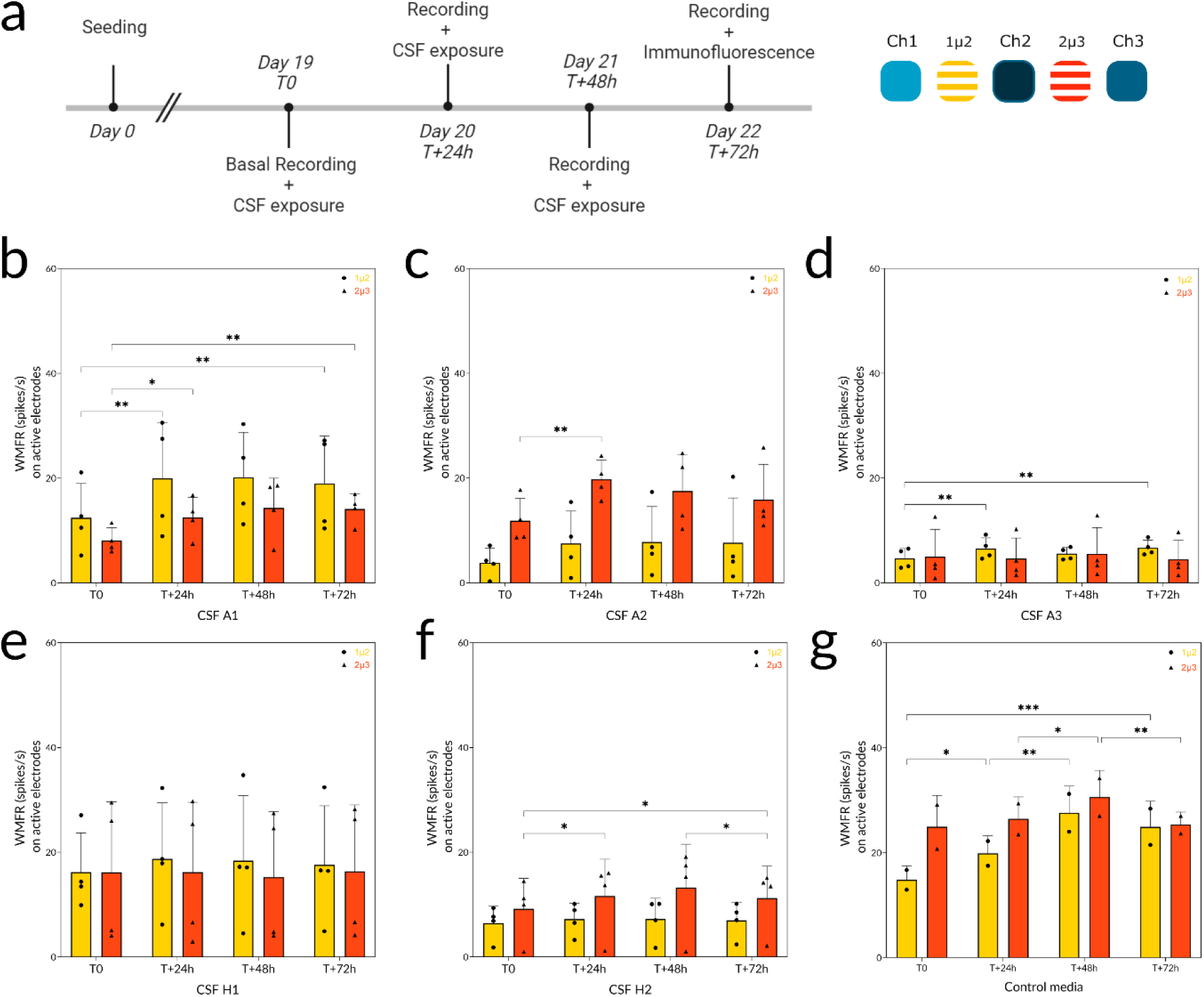
: Electrophysiological effect of AD and healthy patient’s CSF exposure on co-culture glutamatergic neurons. (a) Timeline of CSF exposure experiment steps. (b, c, d) Bar graphs showing WMFR in microchannels of chronic CSF exposure, from three anonymous Alzheimer’s patients called A1, A2, and A3, in time. (e, f) Bar graphs showing WMFR in microchannels of chronic CSF exposure, from two anonymous healthy patients named H1 and H2, in time. (g) Bar graph of WMFR in microchannels of chronic addition of control media in time. * p-value < 0.05, ** <0.01 Error bar = SD. CSF: cerebrospinal fluid.

**Figure 6:**
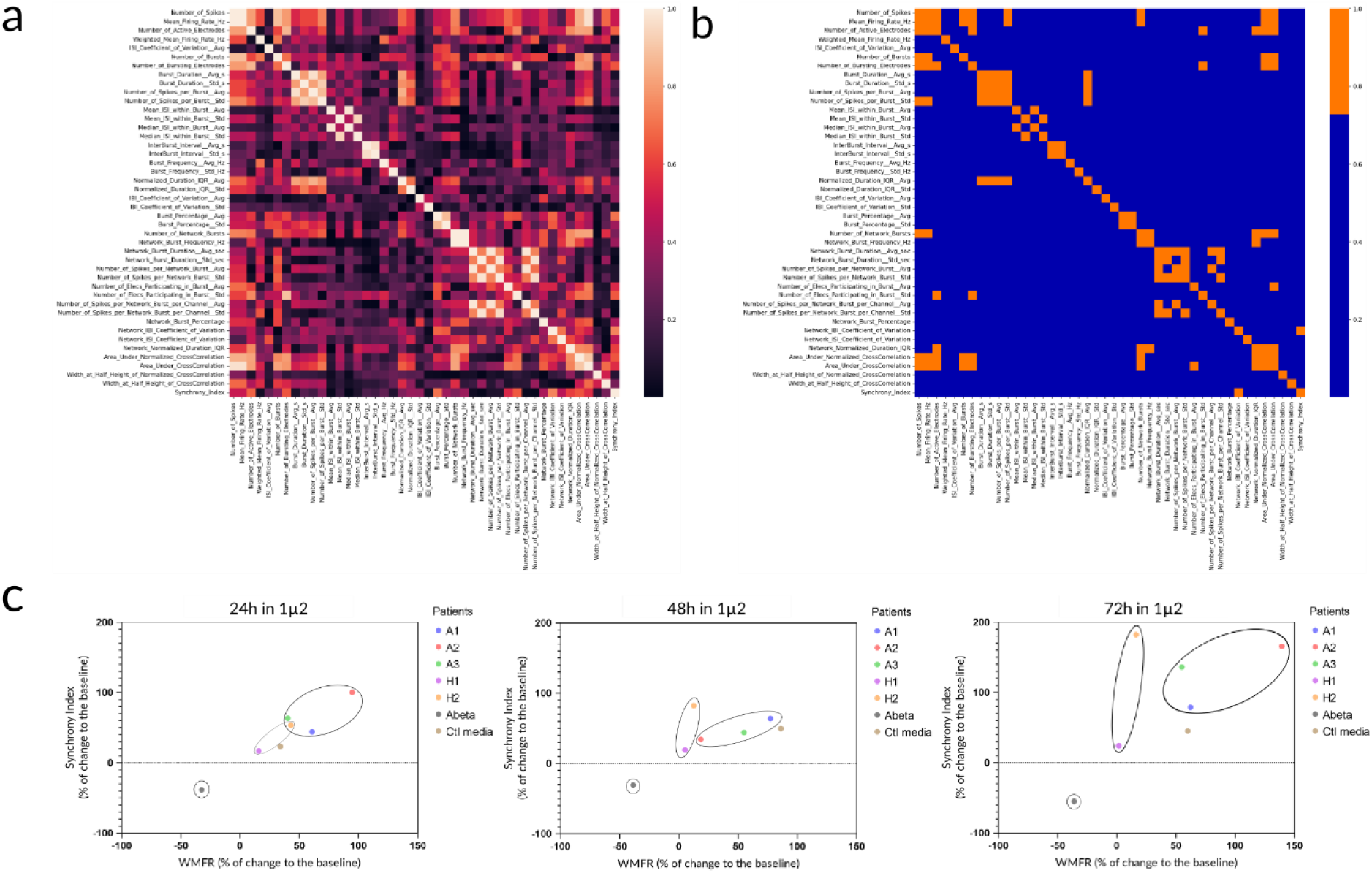
Electrophysiological analysis of CSF from AD and healthy patients’ exposure on glutamatergic neurons. (a) Correlation matrix of normalized dataset from all electrophysiological metrics. Each cells represents the Pearson correlation coefficient between pairs of metrics. Correlation coefficients range from 0 to 1. (b) An arbitrary threshold of 0.7 was applied to differentiate strong (in orange) and weak (in blue) correlations between metrics. (c) Scatter plots of synchrony index and WMFR at 24h, 48h and 72h after experiments in 1µ2 compartment.

## Discussion

In this study, we investigated the effect of AβO and AD patients’ CSF on functional activity of a compartmentalized glutamatergic neurons culture. Prior to expose neurons to AβO and CSF, we ensured that glutamatergic neurons were fully differentiated and capable to respond to chemical stimuli. Recordings in microchannels (1µ2 and 2µ3), have shown a better detection and increase of spikes frequency than in recorded channels (Ch1 and Ch3). This phenomenon, well documented, can be explained by the microchannels architecture^26,27^. Indeed, the confined compartments increase electrode impedance due to the electrical resistance of the microchannel, leading to an increase in spike amplitude thus enhancing signal-noise ratio^28^. Our results demonstrate the inhibitory effect of TTX ^21,29^ on glutamatergic neurons activity, in channels and microchannels, with the decrease of WMFR in TTX-treated channel and axonal compartments. Validation of fluidic isolation, thanks to TTX experiments, allowed us to perform AβO exposure on glutamatergic neurons seeded in Ch1 to observe the effect of neuronal network. We have observed that 10µM of AβO in human glutamatergic neurons have a significant decrease of functional activity after 72h of exposure. However, the used concentration was thousand times higher than Aβ1-42 measured in human CSF from healthy and AD patients^30^ (Supplementary Table S1). Tests using 1µM and 5µM of AβO were also conducted but did not show an effect on functional activity (data not shown). Some research showed that AβO had effect on neurons and synaptic communication at lower concentration as 1, 2 or 5 µM in conventional cell culture as a Petri dish ^31–34^. Several studies have investigated the synaptic effect of Aβ peptides in microfluidic devices made of PDMS, to evaluate the viability^35^, the synapto-toxicity^36^ and the impacted functional activity of neuronal culture^2,37^. Lefebvre et al., in 2024, used an asymmetric microfluidic device allowing to isolate synapses from neuronal culture. They investigated the effect of Aβ on synaptic functional activity in a directional neuronal network showing that applying Aβ to the synapses for 48 hours significantly reduced interchamber connectivity without causing a substantial change in neuronal activity within the presynaptic or postsynaptic chambers^37^. In our study, we used a symmetric microfluidic device to recreate a bi-directional neuronal network, allowing us to observe the effects of Aβ and CSF in glutamatergic neurons (Ch1 and 1µ2, Figure 1d) and assess the impact and the potential response of both cultures (Ch3 and 2µ3, Figure 1d) allowing to mimic the natural interactions that occur in the brain, where neurons communicate in both directions^38^.

AD patients’ CSF exposure on glutamatergic neurons increased the spike rate on active electrodes within 24h and could suggest that AD’s CSF was not cytotoxic to glutamatergic neurons. As previously reported by Yaka et al., CSF of AD patients showed a non-cytotoxicity on neuron-like cells as PC12 ^39^. In the opposite, some publications related the adverse effect of CSF from AD on rodent cell viability and neurotoxic effects ^40,41^. It is known that CSF from neurodegenerative disease could affect *in vitro* glial cells as reported by Schiess et al., with Parkinson’s disease patients’ CSF exposure^42^. It could be explained by the fact that CSF have a cytotoxic effect on in primary cells ^40,41^ rather than cell lines ^39^. In our study, human glutamatergic neurons iPSCs-derived were composed of about 85% of neurons in our microfluidic technology explaining that we didn’t show the same effect of CSF on our culture (Fig. 2h) than on primary cells ^40,41^.

Several studies used healthy and artificial CSF on neural stem cells and have reported that they could promote stem cells proliferation capacity^43^, glial differentiation^43–45^ and neuronal circuit maturation^46^. CSF could have a beneficial effect on human neurons iPSCs derived and then an increase of functional activity, as we shown with CSF from healthy patients (H1 and H2).

In the opposite we expected to have a cytotoxic effect with applying diseased CSF directly on neurons. Several studies reported that exposing CSF from patient suffering from amyotrophic lateral sclerosis on rodent cells and iPSC motor neurons^47,48^ or Parkinson’s disease on dopaminergic neurons^49–51^ have a cytotoxic effect.

Surprisingly, in this study, when we added CSF from patients suffered from AD, we have shown that the functional activity was increased or even improved. In 2013, Grotz et al. ^52^, investigated the effects of AD patients’ CSF from different stage of the disease on functional activity of rodent cortical culture. Similarly to us, they reported an increase in spike rate after AD patients’ CSF exposure compared to baseline. However, they have shown that the activity was decreased with AD patients’ CSF compared to CSF of mild cognitive impairment (MCI) patients, an early stage of AD ^52^. As Görtz et al., we used neuronal media composed of BrainPhys, which is known to have an excitatory effect on neuronal culture. This would explain the increase of functional activity over time in 1µ2 and 2µ3 recorded in the control condition (Fig. 5g). To further our electrophysiological analyses, we performed correlations between two metrics, WMFR and synchrony index. Results expressed in percentage of change to the baseline in scatter plot. At 24h, 48h, and 72h, scatter plots demonstrated a dynamic progression in the differentiation of patient groups (Fig. 6). Initially, at 24h, the groups were relatively undifferentiated. By 48h, emerging trends show a spread and some separation, and by 72h, there is clear isolation and expansion of the groups. This could be due to diverse reactions to treatment, differences in disease progression, or other patient-specific factors. The overall trend indicates a clear stratification of patient groups, which may be crucial for personalized diagnosis approaches or understanding disease mechanisms. A better distinct correlation could be identified with a larger number of samples (AD patients, n=3 and healthy patients n=2). Since Görtz et al.^52^, in 2013, there have been no further publications on the effect of AD’s CSF on neuronal activity using MEA technology. This lack of studies on the subject can be explained by the fact that CSF is a biological fluid whose composition is currently misunderstood. However, Koch et al., in 2019^14^ presented a device and methodology allowing to detect effect of CSF from AE patients in neuronal network activity. They have shown that CSF exposure of AE patients could significantly decrease spike and burst rates.

Investigating the effects of CSF on neuronal cultures *in vitro* presents several challenges due to the complex and multifaceted nature of CSF. It is a dynamic biological fluid that contains a wide array of components, including proteins, metabolites, electrolytes, and signalling molecules as pro-inflammatory and REDOX molecules. Its composition can vary significantly, especially in the context of neurological diseases, and it can be difficult to control the quantity of total components ^53^.

The general approaches that have been used for the last decades to investigate CSF composition were all based on detecting biomarkers and trying to make sense of their ratios. In this paper, we are aiming at developing a novel strategy using a microfluidic platform seeded with human neurons acting as biosensors. This innovative approach allows us to go beyond traditional biomarker detection by harnessing the intrinsic ability of neurons and neural networks to propagate and encode functional information. By exposing the cultured neurons to various biological components in one compartment only, we demonstrated that the targeted neural cell culture is acting as a transducer, while the other connected neural population not exposed to the fluid is acting as a biosensor for which the distinct activity patterns reflect the remote functional impact. This method provides a more dynamic and integrative perspective on how CSF composition influences neuronal function, offering a powerful tool for both research and potential clinical applications in the diagnosis and understanding of neurological diseases.

Despite the difficulty of investigate CSF effect *in vitro*, our system provides reliable and reproducible results, allowing to facilitate the classification of sample and distinguish between different patient subgroups based on their unique biological profiles. Recently, microfluidic technology has been used to investigate biological fluids allowing to develop new tools for AD diagnosis and biomarkers detection ^54–56^. As we shown, microfluidic platform coupled with MEA provides a relevant tool to investigate the functional effect of a perturbation of neuronal culture. However, there are several areas of study that should be explored to improve the model and the sample classification. An oriented unidirectional co-culture could be used to precisely control and analyse synaptic connectivity and signal propagation. They can be created using specific technologies such as axon diodes or arches ^57,58^ to enable only the axons from a specific channel to cross and connect with the other channel, and not the other way around. It is also possible to perform experiments in an asymmetric microfluidic device allowing the synapses isolation and investigate neuronal communication of oriented neuronal network^27,57,59,60^. Then, the improved complexity of co-culture with two complementary neuronal types, such as glutamatergic and GABAergic neurons, would allow to study the excitatory and inhibitory effects and the feedback loops, present in neurological disorders^61–63^. Furthermore, in-depth exploration of all the electrophysiological metrics would increase the sensitivity and specificity of the detection device. Indeed, dozens of metrics can be extracted from the electrophysiological tools used in this study. Visualizing the relationship between metrics, such as scatter plots and cross-correlation graphs could allow us to better distinguish different experimental conditions and treatments. Moreover, correlating metrics can reveal patterns or anomalies that single-metric analyses might miss, leading to more robust and comprehensive conclusions. There is a need to develop automation tools to improve the specificity of detection. Then, the number of patients needs to be significantly increased to obtain reliable responses and increase the sensitivity of the detection platform.

Using neurons as sensors within a microfluidic platform with compartmentalized neurons holds potential for differential diagnosis of neurological disorders. By introducing patient-derived CSF into one compartment, we could monitor the result changes in neuronal activity across the entire network and have a specific pattern for each disease. This approach could be used in the development of personalized therapeutic strategies by assessing how individual patient samples influence neural networks.

## Materials and methods

### High throughput microfluidic devices coupled with microelectrode array (MEA)

Microfluidic devices with three compartments including two symmetrical culture chambers, linked with microchannels were fabricated using Polydimethylsiloxane (PDMS) and conventional photolithography technic as previously described in Honegger et al.6464, called DuaLink MEA (NBP_TLN-AX, NETRI, Lyon). Microfluidic devices were bonded with oxygen plasma in an Axion Biosystem custom-made sheets composed of MEA. MEA devices were filled in with 70% ethanol and three washes of distilled water before coating.

### Neuronal cell culture

#### Rodent cells

Hippocampal and cortical neurons have been collected had previously reported in Maisonneuve et al.6565. Briefly, neurons were sampled from E18 OFA rats (Charles River Laboratories) and digested for 30 min using 2mL of HEPES-buffered HBSS containing 20 U/ml papain (Worthington Biochem., USA), 1mM EDTA (PanReac AppliChem) and 1 mM L-cysteine (Sigma Aldrich, USA). The cells were gently triturated in 1mL of neuronal culture medium, counted with a Malassez cell counter (REF) and flowed into the devices. The cells were maintained under incubation conditions (37 °C, 5% CO2, and 80% humidity) until use.

#### Human cells

Human-induced pluripotent stem cell-derived cortical glutamatergic neurons (BrainXell, BX-0300, Madisson, WI, USA) and GABAergic neurons (BrainXell, BX-0400, Madisson, WI, USA) were purchased. Microfluidic devices were previously coated with PDL (0.05 mg/mL) overnight at 37 ◦C under 5% CO2. Glutamatergic neurons were seeded by placing 3 µL of a 1 × 10^7^ cells/mL neuron suspension in the inlet reservoir of targeted channels, depending on experimental conditions at a density of 600 cells/mm^2^. Cell culture media was composed of several supplement and Brain Phys (Stemcell Technologies, Canada, #05790), known a neuronal activity improvement6666, as providers recommendations. Fresh media were replaced every 2 to 3 days with provider media, as previously described 6767. Neurons were cultured for up to 21 days under a controlled environment (37^°^C and 5% CO2). Human-derived materials were preserved and handled with the approval and under the guidelines of French legislation. The accreditation number related to the use of human materials is DC-2020-4203.

### TTX preparation and exposure

Tetrodotoxin (TTX), a potent and specific blocker of voltage-gated sodium channels has been purchased at Laxotxan (L-8502, Laxotan, Velcen, France) and was prepared at a stock solution of 10mM in sterile phosphate-buffered saline (PBS) and stocked at −20°C. Neurons in culture were exposed to a working solution of TTX diluted in fresh culture media at a concentration of 10nM. Neuronal cultures were exposed to TTX for few minutes. Real-time recordings were taken to record the immediate effects of neuronal firing. Recordings were performed 24h post-TTX exposure on culture to evaluate the recovery of neuronal activity.

### Beta amyloid oligomers (AβO) preparation

AβO were obtained from Etap Lab (Nantes, France). Oligomers were diluted to the targeted concentration and mixed with cell culture media for direct application. As a control, vehicle from AβO preparation buffer and cell culture media have been used. Human amyloid beta 1-42 monomers were obtained from Bachem (Germany). Oligomers were prepared as described previously (Dahlgren et al., 2002). Briefly, amyloid beta peptide was dissolved to 1 mM in 100% hexafluoroisopropanol (HFIP). HFIP was removed under a nitrogen stream, then peptide was resuspended in DMSO to 5 mM. F-12 (without phenol red) culture media was added to bring the peptide to a final concentration of 500 µM, and the peptide was incubated at 22 °C for 18 h. Oligomers were then aliquoted and stored frozen at −80°C until use.

### CSF from patients’ collection and analysis

CSF collection, sampling and storage were performed using a standard procedure according to the international consensus 6868. The concentrations of Aβ1-42, t-Tau, pTau181 and Aβ1-40 were measured routinely using Lumipulse G 600II (Fujirebio) in the neurochemistry laboratory (Hospices Civils de Lyon, Groupement Hospitalier Est, Lyon, France). (Supplementary Table S1). The cut-off values defined by the laboratory, considering a positive AD CSF biomarker profile were t-Tau ≥ 400 ng/L, pTau181 ≥ 60 ng/L, and Aβ1-42 ≤ 550 ng/L and/or a Aβ1-42/ Aβ1-40 ratio <0,055 (Molinuevo et al., 2014). All the assays were performed in the laboratory according to ISO 15189:2012 standard (n°8-3442 rev.28). Patients or their relatives gave their written consent to save CSF for research purposes. Samples were stored in a biobank with authorization from the French Ministry of Health (Declaration number DC-2008-304). Prior to use, CSF was diluted to 10% v/v in cell culture media before addition to culture media. As control, fresh culture media was added on neurons.

### Immunofluorescence

Cultures were fixed in 4% paraformaldehyde (PFA) for 30 min at room temperature, as previously described 6767. Briefly, cells were washed three times with PBS and permeabilized for 10 min with 0.1% Triton-X100, followed by 30 min with 3% BSA. Primary antibodies: mouse anti-Nestin (1:500 (1 µg/mL), ThermoFisher, 14-9843-82), rabbit anti-Sox2 (1:500 (2 µg/mL), Merck, AB5603), guinea pig anti-vGlut1 (1:100 (5 µg/mL), Synaptic Systems, 135304), rabbit anti-GABA (1:50 (5 µg/mL), Sigma-Aldrich, A2052), rabbit anti-MAP2 (1:500 (1 µg/mL), Abcam, ab96378), and mouse anti-βIII-tubulin (1:200 (5 µg/mL), ThermoFisher, MA1-118) were added, and the devices were incubated overnight at 4°C. The cells were rinsed three times with PBS and further incubated with the corresponding secondary antibodies : Alexa Fluor 488 goat anti-mouse IgG (1:1000, Abcam, ab150061), Alexa Fluor 647 donkey anti-rabbit IgG (1:1000, Abcam, ab150075), Alexa Fluor 647 goat anti-guinea pig IgG (1:1000, Abcam, ab150187), Alexa Fluor 555 donkey anti-mouse IgG (1:1000, Abcam, ab150110) and Alexa Fluor 488 donkey anti-rabbit IgG (1:1000, Abcam, ab150061, for 2 h at room temperature in the dark. For the quantification of antibodies, the percentage of the positive cells was quantified compare with DAPI (Sigma-Aldrich, D8417) positive using a homemade macro in Image J. Images were acquired with an inverted epifluorescence microscope, the AxioObserver 7 (Zeiss), with a CMOS camera.

### Live/Dead Assay

Cell viability assessment was performed as previously described 6565. Briefly, cells were incubated with the LIVE/DEAD™ Viability/Cytotoxicity Kit (L3224, Thermo Fisher Scientific Inc., USA), for 30 min at room temperature and were imaged without washing.

### Image analysis

All microscope pictures were treated with a NETRI’s proprietary software developed with ImageJ (National Institutes of Health, Bethesda, MD, USA) allowing to quantify the number of positive cells stained with fluorescence

### Electrophysiological recordings and analysis

Electrophysiological recordings were performed with commercial hardware and software by Axion Biosystems (Axion Biosystems, Atlanta, GA). MEA recordings were performed using the Maestro Pro MEA system (Axion Biosystems). Baseline neuronal activity was recorded for 10 minutes before any treatment use as a control baseline of activity. The recorded data were analyzed using AxIS Navigator software (Axion Biosystems) and NETRI’s proprietary software NETRI UpLink, which facilitated the extraction of electrophysiological metrics such as MFR and raster plots, WMFR and synchrony index, for each compartment (Fig 1f).

The threshold for spike detection was set at 6x the standard deviation of the noise level of the root mean square (RMS), which was determined from the initial segment of the recording without evident neuronal activity. This method ensured reliable discrimination of true neuronal spikes from background noise.

The mean firing rate (MFR) is the number of spikes per second from each channel and was calculated on all the electrodes divided by recording time, while the weighted mean firing rate (WMFR) corresponds to the number of spikes per second from each channel and was calculated only on the signal detected on the active electrode. An active electrode has been considered when the electrode detects at least 0.1 spikes/s. Recordings with an activity less than 0.1 spikes per second (spikes/s) at day 18 (D18) for TTX experiments and at day 20 (D20) for other tests (including Aβ and CSF treatments) were excluded from the analyses.

## Data Analysis

Data were analyzed using GraphPad Prism version 10.2.2 (GraphPad Software, San Diego, CA, USA). All experimental results are presented as mean ± standard deviation (SD) or Standard Error of the Mean (SEM) for n>10. Data points from individual experiments are shown alongside the mean values to illustrate variability. For comparisons between multiple groups (days of experiment), a two-way analysis of variance (ANOVA) was performed. Post-hoc multiple comparison tests were conducted to identify significant differences between groups using Tukey’s test or Fisher’s LSD test depending on analyzed samples. The significance level was set at p < 0.05. Graphs were generated using GraphPad Prism. Statistical significance in the graphs is indicated by asterisks (*p < 0.05, **p < 0.01, ***p < 0.001 and ****p < 0.0001).

## Acknowledgements

The authors received no specific funding for this work.

## Author contributions

Conceptualization, L.M., T.H. and I.Q.; experiments, L.M., L.D., A.B., F.B. and M.H..; electrophysiological analyses, L.M.; data analyses: L.M.; A.P.; validation, L.M., J.R., F.L., I.Q. and T.H.; writing, L.M..; review, T.H., I.Q.

