## Supplementary material for "Neurons as biosensors for discriminating neurological disorders in a brain-on-chip platform: Application to Alzheimer’s Disease using patient CSF": Supp figures

| Patients ID | Age at sampling | Diagnostic | Tau total (pg/l) | Tau P181 (ng/l) | Aβ 1-42 (ng/l) |
|---|---|---|---|---|---|
| A1 | 60 | AD | 585 | 105 | 408 |
| A2 | 78 | AD | 523 | 65 | 491 |
| A3 | 82 | AD | 876 | 174 | 496 |
| H1 | 75 | Healthy | 205 | 33 | 1051 |
| H2 | 72 | Healthy | 335 | 46 | 726 |

Table S1: Clinical information of selected patients' CSF samples. AD: Alzheimer's disease.

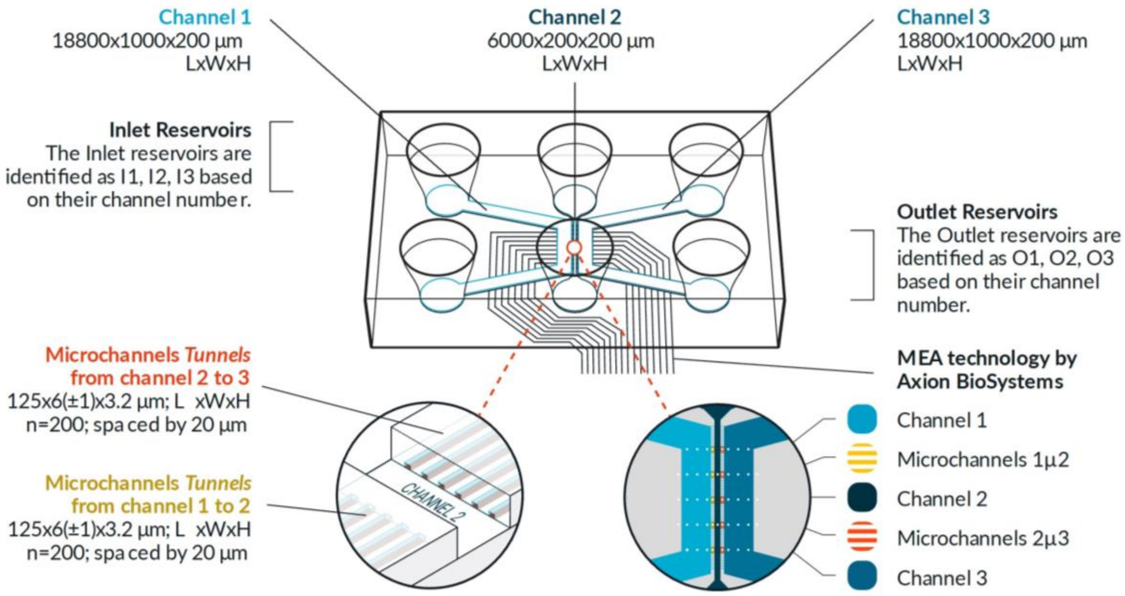


Figure S1: Schematic representation of DuaLink MEA with dimensions.

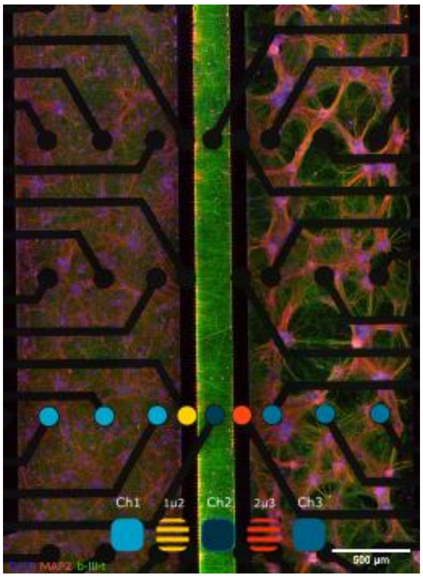


Figure S2: Immunofluorescence picture of rodent co-culture of cortical and hippocampal neurons seeded in the Dualink MEA. Neurons were stained with B-III-tubulin in green, MAP2 in red, DAPI in blue. The third row of electrodes has been coloured to identify electrodes below each compartment.

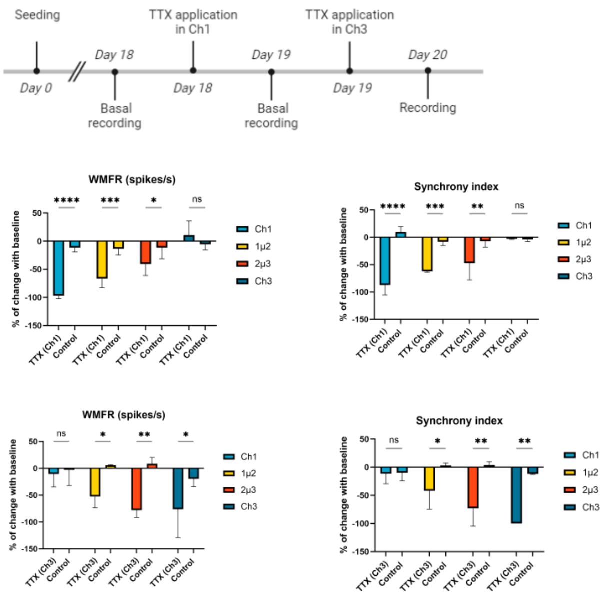


Figure S3: Electrophysiological modulation of co-culture glutamatergic neurons with TTX and vehicle. (a) Timeline of TTX and vehicle exposure experiment steps. Bar graphs showing WMFR and synchrony index overtime in channels and in microchannels respectively, expressed in percentage of change with baseline. * p-value < 0.05, ** p-value <0.01, *** p-value < 0.001 and **** p-value < 0.001. Error bar = SD. TTX: tetrodotoxin. WMFR: weighted mean firing rate

### Coomassie blue stained SDS-PAGE of AβO:

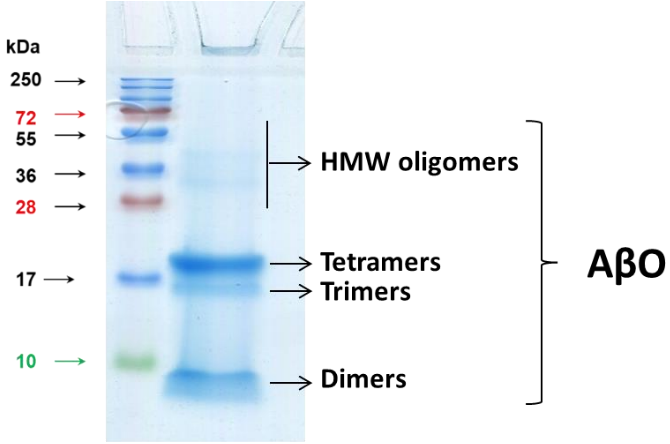


Figure S4: AβO were analyzed using 15% SDS-PAGE gel and Coomassie-blue staining. Gel profile demonstrated that oligomer preparations contain a mixture of dimers, trimers, tetramers, as well as traces of high-molecular weight Aβ oligomers. Molecular weight markers were loaded in the first lane, and the molecular masses are indicated.

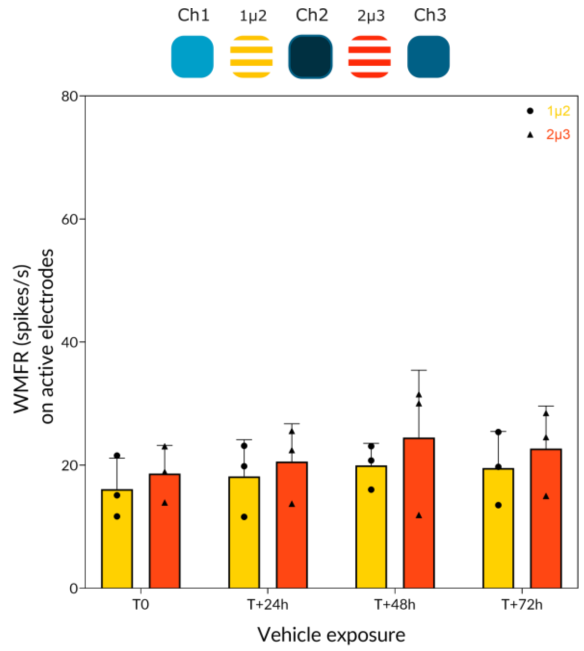


Figure S5: Electrophysiological effect of Aβ vehicule exposure on co-culture glutamatergic neurons. (a) Timeline of Aβ vehicle exposure experiment steps. Bar graph of WMFR in microchannels of chronic vehicle exposure in time respectively. Error bar = SD. WMFR: weighted mean firing rate.
